# Multi-spectral photoacoustic imaging combined with acoustic radiation force impulse imaging for applications in tissue engineering

**DOI:** 10.1101/2024.04.23.590806

**Authors:** Christopher D. Nguyen, Ying Chen, David L. Kaplan, Srivalleesha Mallidi

## Abstract

Tissue engineering is a dynamic field focusing on the creation of advanced scaffolds for tissue and organ regeneration. These scaffolds are customized to their specific applications and are often designed to be complex, large structures to mimic tissues and organs. This study addresses the critical challenge of effectively characterizing these thick, optically opaque scaffolds that traditional imaging methods fail to fully image due to their optical limitations. We introduce a novel multi-modal imaging approach combining ultrasound, photoacoustic, and acoustic radiation force impulse imaging. This combination leverages its acoustic-based detection to overcome the limitations posed by optical imaging techniques. Ultrasound imaging is employed to monitor the scaffold structure, photoacoustic imaging is employed to monitor cell proliferation, and acoustic radiation force impulse imaging is employed to evaluate the homogeneity of scaffold stiffness. We applied this integrated imaging system to analyze melanoma cell growth within silk fibroin protein scaffolds with varying pore sizes and therefore stiffness over different cell incubation periods. Among various materials, silk fibroin was chosen for its unique combination of features including biocompatibility, tunable mechanical properties, and structural porosity which supports extensive cell proliferation. The results provide a detailed mesoscale view of the scaffolds’ internal structure, including cell penetration depth and biomechanical properties. Our findings demonstrate that the developed multimodal imaging technique offers comprehensive insights into the physical and biological dynamics of tissue-engineered scaffolds. As the field of tissue engineering continues to advance, the importance of non-ionizing and non-invasive imaging systems becomes increasingly evident, and by facilitating a deeper understanding and better characterization of scaffold architectures, such imaging systems are pivotal in driving the success of future tissue-engineering solutions.

## 1. Introduction

Traditional methods of repairing damaged tissues and organs involve grafts or transplants, but issues such as immune rejection and limited donor supply often hinder these options. Among the array of alternative approaches, engineered tissues utilizing scaffolds infused with patient-derived tissues and cells stand out as an excellent candidate. When designing these scaffolds, several factors including biocompatibility, biodegradability, mechanical properties, and structure must be considered. While an abundance of materials has been studied for this purpose, one of notable interest is silk fibroin protein (silk hereafter) derived from Bombyx mori silkworms. Silk is an established material in the field of biomaterials and tissue engineering due to its desirable properties of the aforementioned factors, and its ability to form interconnected porous structures which allow cell proliferation into the scaffold [1]. Furthermore, these scaffolds can be fabricated into various shapes and sizes with complex geometries that mimic the native tissue environment to ensure proper cell behavior [2]. Current imaging methodologies to examine these intricate scaffolds focus on microscale evaluation such as scanning electron microscope (SEM) and micro-CT to view micropores, or optical imaging techniques such as the confocal or fluorescence to examine cell composition and migration [3, 4]. However, due to their limited penetration depth, these techniques are inadequate when the scaffolds reach millimeters or centimeters in size. Additionally, the capability of measuring the scaffold stiffness which has been shown to affect cellular adhesion and proliferation is a critical feature [5, 6]. Therefore, as the tissue engineering field is advancing towards developing larger artificial tissues, a multi-parametric imaging system is desired which can provide complementary 3D structural, compositional, and mechanical properties at ample depths and sufficient resolution.

Optical imaging techniques utilized to image tissue engineered constructs tend to rely on optical focusing to provide high resolution, but at the cost of penetration depth due to tissue scattering. While there have been several advances in optical microscopy techniques to probe deeper into tissues such as with recent advances in light sheet microscopy [7-9], they are still limited to depths of less than 500 μm. Conversely, ultrasound (US) imaging techniques can image thicker tissue engineered constructs to provide unprecedented structural information, but at lower resolution. However, the frequency of the ultrasound transducer can be changed according to the desired application to trade off between resolution and penetration depth. To obtain compositional and functional data at ample penetration depths and sufficient resolution while leveraging the advantages of optical contrast, ultrasound based photoacoustic (PA) imaging that relies on the photoacoustic effect to convert optical to acoustic energy may be used. PA imaging traditionally utilizes multi-wavelength sources to capture spectral data which is unmixed to provide the compositional and functional data, most notably highly absorbing contrast agents or blood oxygenation [10-15]. PA is commonly combined with US due to their similar receiver electronics and complementary data sets. In a similar manner, acoustic radiation force impulse (ARFI) imaging may be combined with ultrasound to obtain complementary mechanical properties [16-18]. ARFI delivers an intense focused acoustic pulse to the sample to induce a local displacement and relaxation which is tracked using regular pulse-echo ultrasound. The amplitude of the displacement aids in gauging the sample stiffness.

Previously, US and ARFI have been combined with either fluorescence [19] or shear wave elasticity imaging (SWEI) [20] to obtain additional information on the tissue samples. Multi-spectral fluorescence imaging has been shown to differentiate malignant from normal tissue due to their differing auto-fluorescence emission spectrum, while SWEI has been shown to provide quantitative characterization of sample stiffness. However, fluorescence is not fully complementary with US and ARFI due to its limited surface penetration, and SWEI in a microscopy configuration has less spatial resolution than ARFI. In a similar study, photoacoustic imaging has likewise been exclusively combined with ARFI in a microscope configuration, however, the system utilized PA to detect sample displacement rather than as a separate imaging modality [21].

To address the aforementioned needs in 3D imaging of millimeter-scale tissue engineered samples at sufficient resolution, we propose the combination of US, PA, and ARFI into a single system to capture spatially co-registered meso-scale information of tissue engineered constructs. Based on our desired scanning volume and resolution, the system was designed in a microscope configuration using two co-linearly aligned transducers and a concentrically aligned optical fiber bundle. This system combining the three synergistic imaging modalities, referred henceforth to as M-PAUR (Multi-spectral PhotoAcoustic, Ultrasound, and acoustic Radiation force impulse), was evaluated using carbon fibers for ultrasound and photoacoustic resolution, and gelatin phantoms for ARFI resolution. To validate its use in imaging tissue engineered scaffolds, M-PAUR was used to confirm the constructed scaffold geometries, incorporated materials, and mechanical properties of silk fibroin-based scaffolds with and without cells. Success in displaying the macro structure, photoacoustic contrast, and stiffness of the scaffolds, in addition to displaying cell depth proliferation as a function of silk concentration, demonstrated the use of M-PAUR in evaluating engineered scaffolds.

## 2. Methods and Materials

### 2.1. Preparation of gelatin phantoms

Gelatin phantoms were prepared by mixing gelatin powder (G2500, Sigma-Aldrich) and titanium oxide (232033, Sigma-Aldrich) into deionized (DI) water. The DI water was heated to 40 °C using a hotplate and gelatin powder (4% w/w and 10% w/w) was slowly added and stirred using a magnetic stir rod. Once the gelatin powder was fully dissolved, titanium oxide (3% w/w) was slowly added and stirred for 20 minutes. The solution was then cooled to 30 °C and poured into rectangular molds (50 mm x 50 mm x 4 mm). For phantoms used to test the ARFI lateral resolution, one concentration was poured into the square mold to approximately 4 mm in height and once solidified, was cut in half and a piece was removed. The second concentration was then poured into the empty space and allowed to solidify. For phantoms used to test the ARFI axial resolution, one concentration was poured into the square mold to approximately 2 mm in height and once solidified, the second concentration was poured directly on top to approximately 2 mm in height and allowed to solidify.

### 2.2. Preparation of silk phantoms

Silk phantoms were prepared using *B. mori* silkworm cocoons that were first cut into small pieces and boiled in a sodium carbonate solution to degum the fibers. The resulting fibers were rinsed to remove the sodium carbonate solution and dried overnight to remove excess water. These dried fibers were added to a 9.3 M lithium bromide solution and incubated to dissolve the fibers into solution which was added to dialysis cassettes for dialysis in ultrapure water. The resulting solution of silk protein in water was centrifuged to remove impurities and poured into a 35 mm petri dish to be lyophilized and autoclaved after adjusting the concentration to 4% or 8%. Further in-depth explanations of the extraction and preparation process can be found elsewhere [22].

The two-part silk scaffold consisted of an 8% silk disk embedded into a 4% silk annulus. To provide photoacoustic contrast, graphite powder (282863, Sigma-Aldrich) was incorporated into the 8% silk solution before lyophilization. To assemble the disk and annulus, the autoclaved 35 mm silk disks were rehydrated in water and shaped using different diameter biopsy punches. The 8% silk disk had a diameter of 3 mm while the 4% silk annulus had an outer diameter of 5 mm and an inner diameter of 3 mm. For scaffolds used to seed melanoma cells, the 35 mm disks were cut into 5 mm diameter disks.

### 2.3. Preparation and seeding of murine melanoma onto silk scaffolds for paraffin sectioning and staining

Melanoma cells (B16-F10, ATCC) were grown and sub-cultured using Dulbecco’s Modified Eagle’s Medium 1x (Corning) supplemented with 10% fetal bovine serum (Gibco). In preparation for scaffold seeding, cells were extracted using 0.05% trypsin-EDTA solution (Corning), centrifuged, and counted to create a suspension of concentration 250,000 cells/mL. To determine the required seeding volume, individual silk phantom cylinders were dehydrated using a vacuum line and slowly rehydrated by depositing a known volume of media using a micropipette until the scaffold was fully saturated. Scaffolds were then fully immersed in media overnight to increase cellular adhesion. During seeding, scaffolds were again dehydrated and seeded with the suspended cell solution using the previously measured seeding volume. After 1 hour in a 37 °C incubator, seeded silk scaffolds were completely immersed in media and returned to the incubator. Media was changed every other day until the desired growth period was reached.

After imaging, scaffolds were embedded in paraffin for sectioning and staining. Paraffin embedding followed established protocols using a standard wash of formalin, ethanol, and xylene before immersing in melted paraffin and solidifying overnight in a 4 °C fridge. To stain, scaffolds were sectioned into 60 μm thick slices using a microtome and then dehydrated overnight on a 37 °C hot plate. Upon deparaffinization and rehydration through a standard xylene and ethanol wash, sectioned scaffolds were stained with hematoxylin and eosin (H&E) using an established protocol.

### 2.4. System design of multimodal microscope

The developed multimodal microscope incorporated an optical fiber bundle and two transducers to deliver the optical and acoustic energy required for photoacoustic, ultrasound, and ARFI imaging as shown in Figure 1a. For photoacoustic imaging, an OPO (Phocus HE Benchtop, OPOTEK) was collimated using an achromatic doublet (AC254-030-B-ML, Thorlabs) and the subsequent beam was reflected off a fold mirror (BB2-E03, Thorlabs) into a power meter acting as a beam dump for one minute to allow energy stabilization. Once stabilized, the fold mirror was lifted out of the optical path and the collimated beam was sampled using the aforementioned power meter and a pellicle (PBS-2, Newport) to monitor the OPO laser fluctuation. The unsampled portion of the beam was focused into a fiber bundle using an aspheric condenser (ACL2520U-B, Thorlabs). The bundle consisted of seven polished multimode fibers (FT1000EMT, Thorlabs) epoxied together towards the proximal end and inserted into a metal tube (96990A903, McMaster-Carr) for mounting. The fanned distal end was attached to a custom 3D-printed transducer-and-fiber mount which was configured to mimic an optical condenser. Upon energy absorption by the sample, the generated photoacoustic signal was detected using a 1-inch focused 20 MHz transducer connected to a pulser/receiver (DPR500, JSR Ultrasonics) for amplification. The amplified signal was digitized by a data acquisition card (CSE161G2, GaGe) with a sampling frequency set to 250 MHz. After 55 μs to fully collect the PA signal, the pulser/receiver generated an US excitation pulse to produce an US echo. For digitization, the same pipeline as PA was used.

**Figure 1.**
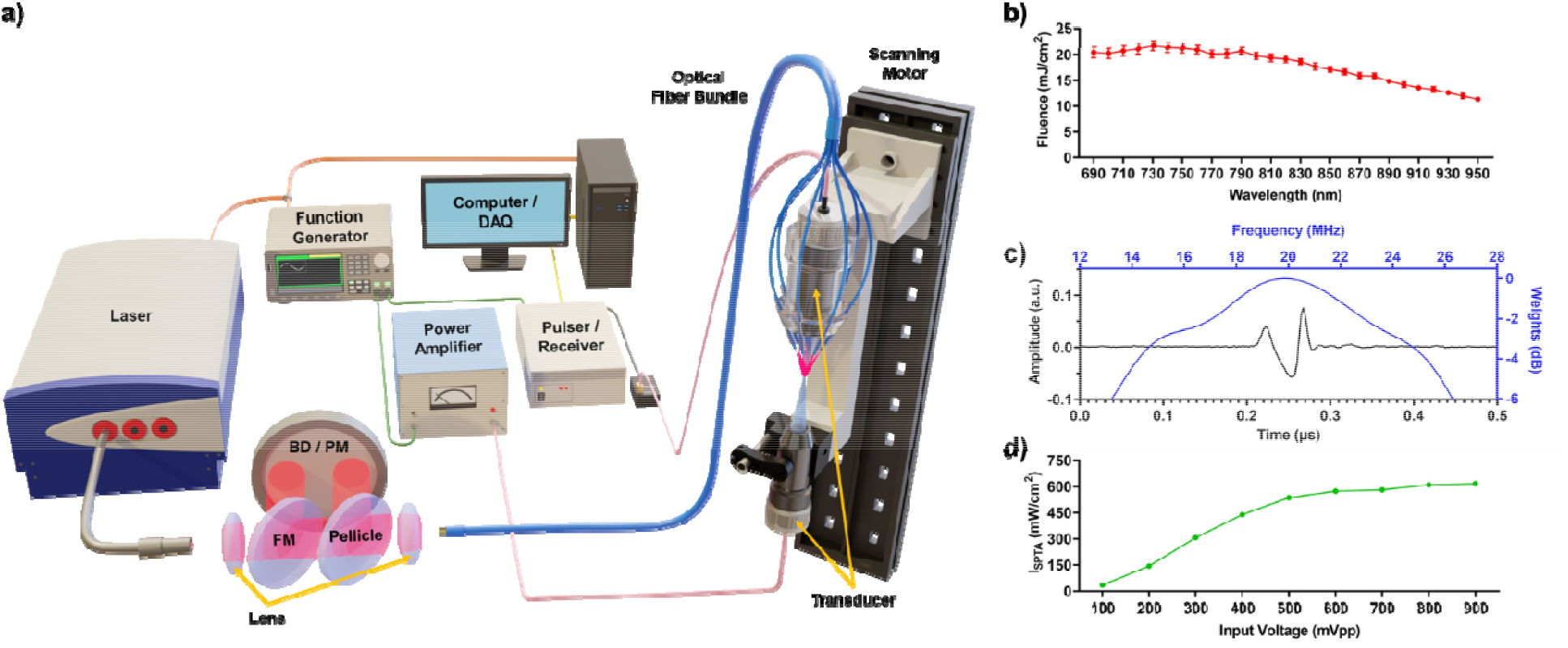
**a)** Schematic of M-PAUR. Laser output is coupled into an optical fiber bundle which splits into an optical condenser configuration. BD: beam dump, PM: power meter, FM: flip mirror. **b)** Laser fluence through optical fiber bundle as a function of wavelength. **c)** Bandwidth of 20 MHz imaging transducer (blue) btained from hydrophone measurement (black). **d)** Generated I_SPTA_ of 5 MHz pushing transducer as a function of input voltage.

To integrate ARFI with PA and US, a 1-inch focused 5 MHz transducer (A310S-SU, Olympus) used for sample displacement was installed on the aforementioned 3D-printed mount and aligned to the 20 MHz transducer by maximizing the US signal obtained when probing a carbon fiber. To generate the acoustic radiation force, a sinusoidal burst sequence from a function generator (DG4102, RIGOL) was amplified by a power amplifier (A-150, ENI) and subsequently passed to the 5 MHz transducer. The sinusoid frequency was set to 5 MHz to match the central frequency of the pushing transducer and the burst cycle was set to 1000. Measurement of the generated pressure and spatial-peak-temporal-average intensity (I_SPTA_) was performed using a hydrophone (HGL-0085, ONDA). To begin ARFI imaging, a reference signal was collected using regular US. An ARFI pulse was then delivered to the sample using the 5 MHz transducer and the resulting sample displacement was tracked with the 20 MHz transducer using 50 US tracking signals triggered at 20 kHz.

Due to the co-linear alignment of the two transducers, the sample was placed on a transparent film (Tegaderm, 3M) to position it at the shared focus. To perform 2D and 3D imaging, the microscope was raster-scanned across the sample and additionally in depth for ARFI. Movement was performed using a 3D motorized stage built using three linear stages (X-LSM, Zaber). Control of the motors to define the imaging volume and step size in addition to the management of data storage was performed using a custom LabVIEW graphical user interface. Further off-line processing of the collected data to generate images was performed using custom MATLAB codes.

Upon completion of building M-PAUR, common microscope parameters were measured. For photoacoustic imaging, the peak fluence output of the laser through the optical fiber bundle was set and confirmed to produce a fluence of approximately 20 mJ/cm^2^ to conform to ANZI standards as shown in Figure 1b [23]. The 20 MHz transducer which collects the resulting photoacoustic signal and performs US imaging was measured to have an approximate bandwidth of 65% or 12.9 MHz (Figure 1c). For ARFI, when the burst cycle was kept constant at 1000, the measured I_SPTA_ spanned between 35 mW/cm^2^ - 620 mW/cm^2^ when the input voltage was increased from 100 mVpp to 900 mVpp (Figure 1d).

### 2.5. Data analysis

The I_SPTA_ was calculated following Equation 1 which relies on the derated pulse intensity integral (PII_derated_) and the pulse repetition frequency (PRF) of the transducer during acquisition [24]. The non-derated PII was acquired by calculating the integral of the pressure waveform squared between and (time interval in which the hydrophone signal is non-zero) and multiplying it by the product inverse between (density of water) and (speed of sound of water). The pressure waveform was acquired using a hydrophone and the manufacturer provided sensitivity conversion from volt to pascal. The speed of sound was assumed to be with the temperature of water at 23.7 °C [25]. Attenuation of the PII from the scaffold followed an exponential decay which assumes for 4% silk and relies on (transducer central frequency) and (distance between transducer and depth of interest) [26].

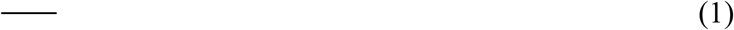

To generate PA and US images, a Hilbert transform was applied to the acquired signals to obtain the signal envelope. To reduce image pixelation, the 3D data set was linearly up-interpolated by a factor of 3 in both the X and Y direction. To display PA images, a simple hot colormap was applied with a manually set amplitude range that provided the best visual contrast. To display US images, the data was modified from a linear to a logarithmic scale that was manually set between a negative value and 0 to provide the best visual contrast. The color was set to a simple gray-based colormap. Formation of an ARFI image involves a cross-correlation between the reference RF signal and the tracking RF signals. Both were segmented to a window size of 1.5 centered between the shared focus of both transducers [27]. However, before cross-correlation, all windowed RF signals were up-interpolated by a factor of 10 using a cubic-spline interpolation [19]. The maximum of the cross-correlation was set as the displacement value for the current voxel. For display, the 3D data set was passed through a 3×3×3 pixel median filter for smoothing and then linearly up-interpolated by a factor of 5 in all directions.

To determine the lateral and axial resolution of PA and US, a carbon fiber acting as a point object was imaged and the full-width-half-max (FWHM) of the point spread in the respective directions was considered the resolution. To determine the lateral and axial resolution of ARFI, gelatin phantoms of two concentrations with transition edges in the lateral and axial directions, respectively, were imaged. A sigmoid function was fit to the measured edge spread to obtain the edge spread function (ESF) and the derivative of the ESF provided the line spread function (LSF). The FWHM of the LSF was considered the resolution.

## 3. Results

### 3.1. Performance of M-PAUR

When imaging the carbon fiber to determine the PA and US resolution, the step size was set to 10 μm laterally as it was sufficiently smaller than the theoretical resolution achievable by the 20 MHz transducer. Based on the FWHM of the point spread, the measured lateral and axial resolution was 344 μm and 55 μm, respectively, for PA and 278 μm and 54 μm, respectively, for US as shown in Figure 2a-c in which US was displayed in grayscale and PA was displayed in a hot colormap. When imaging the gelatin phantoms to determine ARFI resolution, the step size was set to 25 μm laterally and in depth, and the utilized burst sequence was set at 1000 cycles with an amplitude of 500 mVpp which is at the edge of the amplifier’s linear power regime. Assuming a PRF of 10 Hz, this generated an I_SPTA_ of 536 mW/cm^2^ which was more than sufficient to locally displace gelatin phantoms. Based on the resulting FWHM of the LSF calculated from the sigmoid fit of the displacement, the measured lateral and axial resolution was 581 μm and 459 μm, respectively, as shown in Figure 2d-i in which ARFI was displayed in a jet colormap. To satisfy the Nyquist criteria according to these measurements, subsequent PA and US images were obtained with a step size of 100 μm laterally while subsequent ARFI images were obtained with a step size of 200 μm laterally and in depth.

**Figure 2.**
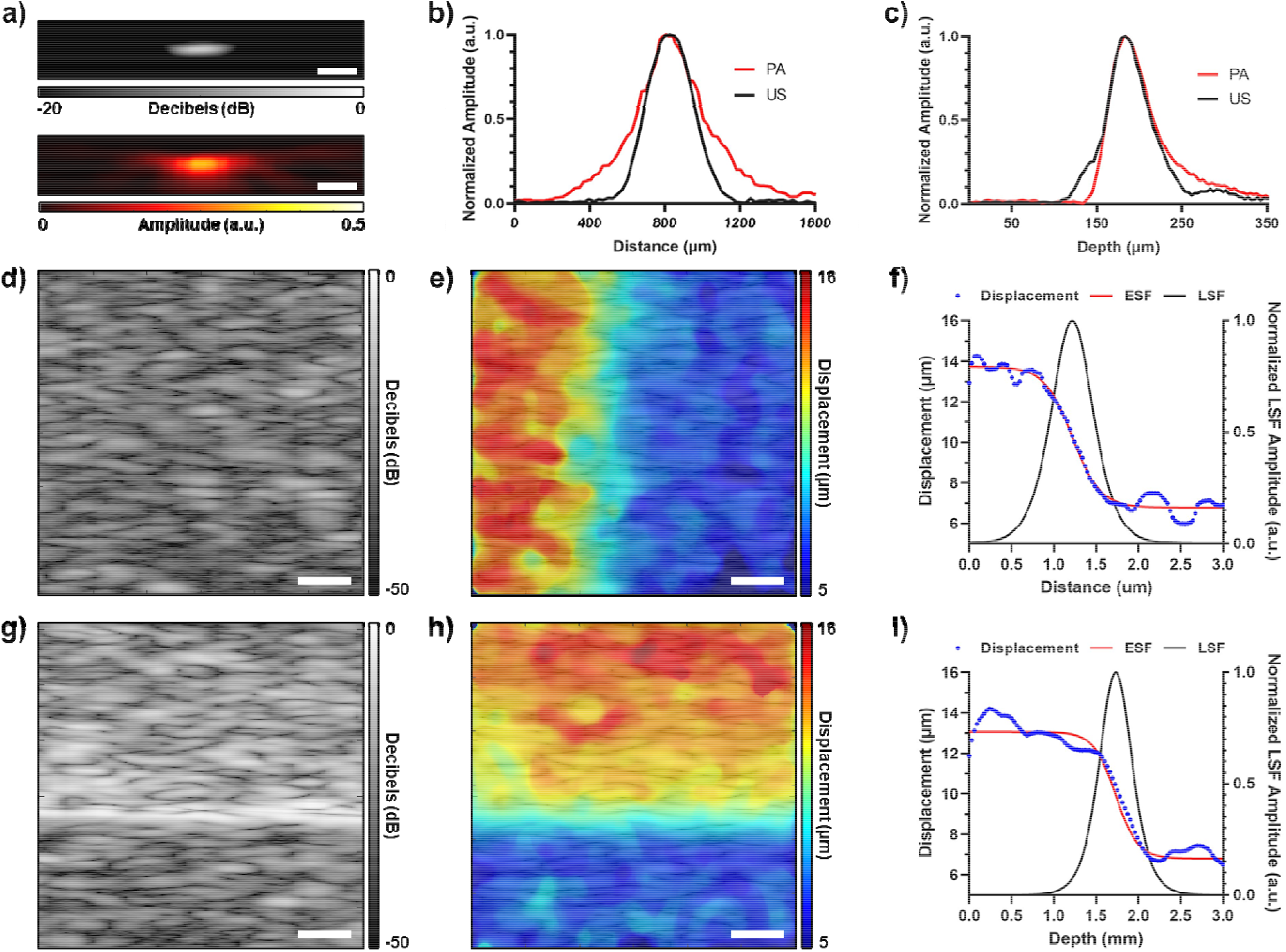
Performance of M-PAUR. **a)** US (top) and PA (bottom) images of a carbon fiber. Scale bar = 250 μm. **b-c)** Lateral and axial profile of carbon fiber images. **d-e)** US and ARFI overlayed image of gelatin phantom with a lateral boundary. Scale bar = 500 μm. **f)** ESF and LSF of lateral boundary from e). **g-h)** US and ARFI overlayed image of gelatin phantom with an axial boundary. Scale bar 500 = μm. **i)** ESF and LSF of axial boundary from h).

### 3.2. Confirmation of detecting contrast in two-part scaffolds

Using the aforementioned step sizes, silk disk scaffolds of two concentrations described in section 2.2 were imaged in 3D for verification of M-PAUR performance. For PA imaging, the wavelength was 690 nm and for ARFI imaging, the burst sequence was set at 1000 cycles with an amplitude of 500 mVpp. Shown in Figure 3, spatially co-registered PA and ARFI images were overlayed onto US. Similar to Figure 2, US images are displayed in grayscale, PA images are displayed in a hot colormap, and ARFI images were displayed in a jet colormap. In Figure 3a-c, pixels located in the colored boxed areas were compared to display differences in acoustic echo, optical absorption, and mechanical properties. Near the top of the phantom, ultrasound signals showed little statistical difference in amplitude between the center and side, however, simple morphological features can be observed such an increased height on right side of the annulus (Figure 3a and 3e). In patially corresponding photoacoustic signals, the center disk with graphite powder showed an approximately 10x stronger signal amplitude compared to the surrounding annulus lacking graphite powder (Figure 3b and 3f). In spatially corresponding ARFI signals, the center disk experienced an approximately 3x lower displacement from the ARFI pulse compared to the surrounding annulus (Figure 3c and 3g). Co-localized images of the three modalities are shown in Figure 3d and Figure 3h depicting overlays in the same plane as Figure 3a-c and en face, respectively. The difference in signal intensity for PA was expected due to the overwhelmingly strong absorption of graphite powder compared to silk which we have previously reported to have an optical absorption coefficient of 0.6 cm^-1^ [26]. Similarly, the difference in signal intensity of ARFI was expected due to the increased material density with higher silk concentration leading to an increased stiffness. These differences were confirmed to be statistically significant upon performing an unpaired two-tailed Mann-Whitney test with a 95% confidence level.

**Figure 3.**
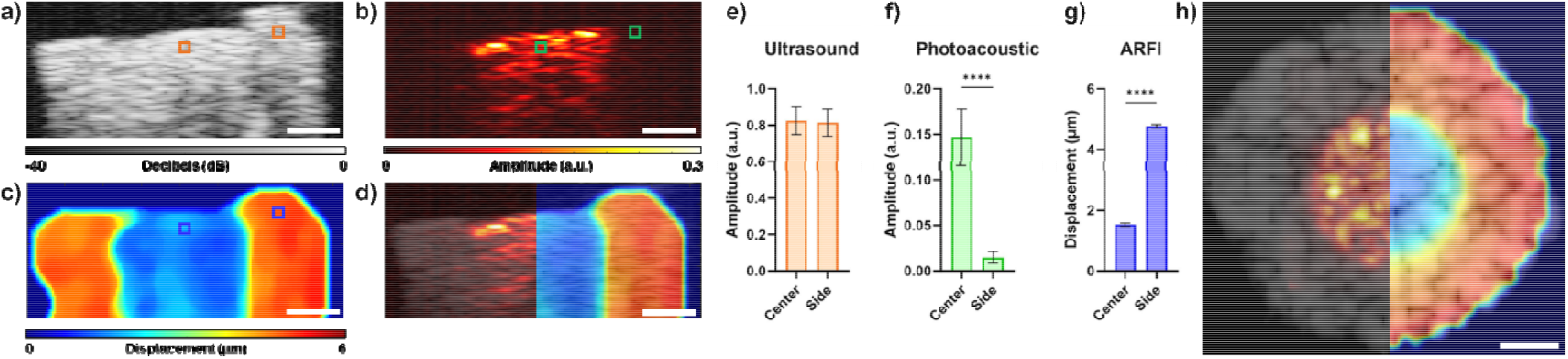
Two-part silk phantom imaged with all M-PAUR modalities. **a-c)** US, PA, and ARFI B-mode images, respectively. **d, h)** Overlayed cross sections of all imaging modalities in the scanning plane and en face,respectively. **e-g)** Bar graphs of pixel amplitudes within a-c), respectively. ****p < 0.0001. All error bars = s.d. All scale bars = 1 mm.

### 3.3. Tracking proliferation of cells within the scaffold as a function of scaffold stiffness

Silk disks seeded with melanoma cells were imaged to measure cell proliferation as shown in Figure 4 following similar imaging parameters used for verifying M-PAUR performance. Melanoma cells were used as the endogenous melanin content generates high photoacoustic signal amplitude due to high optical absorption. For comparison, silk scaffolds prior to cell seeding (labeled as “Day 0”) were imaged to show the ab ence of PA signal originating from the scaffold. In Figure 4 row 1, photos of silk scaffolds are shown with comparative H&E stains in row 2. On day 0, scaffolds appear milky white without cells and expectedly become darker on progressive days as melanoma cells proliferate as shown in H&E stains in which observed black dots correspond to the melanoma cells. Magnified insets of the H&E stains are shown in Fig. S1 which showcases the presence of melanoma cells in the scaffolds at various depths. Row 3 displays ultrasound images which match the macro structure of scaffolds displayed in the photos and H&E stains. The expected speckle pattern has been described in our previous paper which attributes the speckle to the porous alveoli structures generating multiple scattering events [26]. Additionally described in our previous paper, silk exhibits low photoacoustic signal due to its relatively low absorption coefficient as shown in row 4 day 0 photoacoustic images. After 4 days of incubation for both 4% and 8% silk scaffolds, PA images show melanoma cells accumulated towards the top of the scaffold at the site of seeding. By day 8 of incubation, cells are seen proliferating deeper into both scaffold concentrations, however, 4% silk displayed further invasion compared to the 8% silk due to the lower density of pores providing less resistance to cell proliferation. The density of pores can be correlated to scaffold stiffness which is represented in row 5 ARFI images. As previously mentioned, higher silk concentration results in increased material density and therefore increased stiffness which experiences less displacement in response to an ARFI pulse.

**Figure 4.**
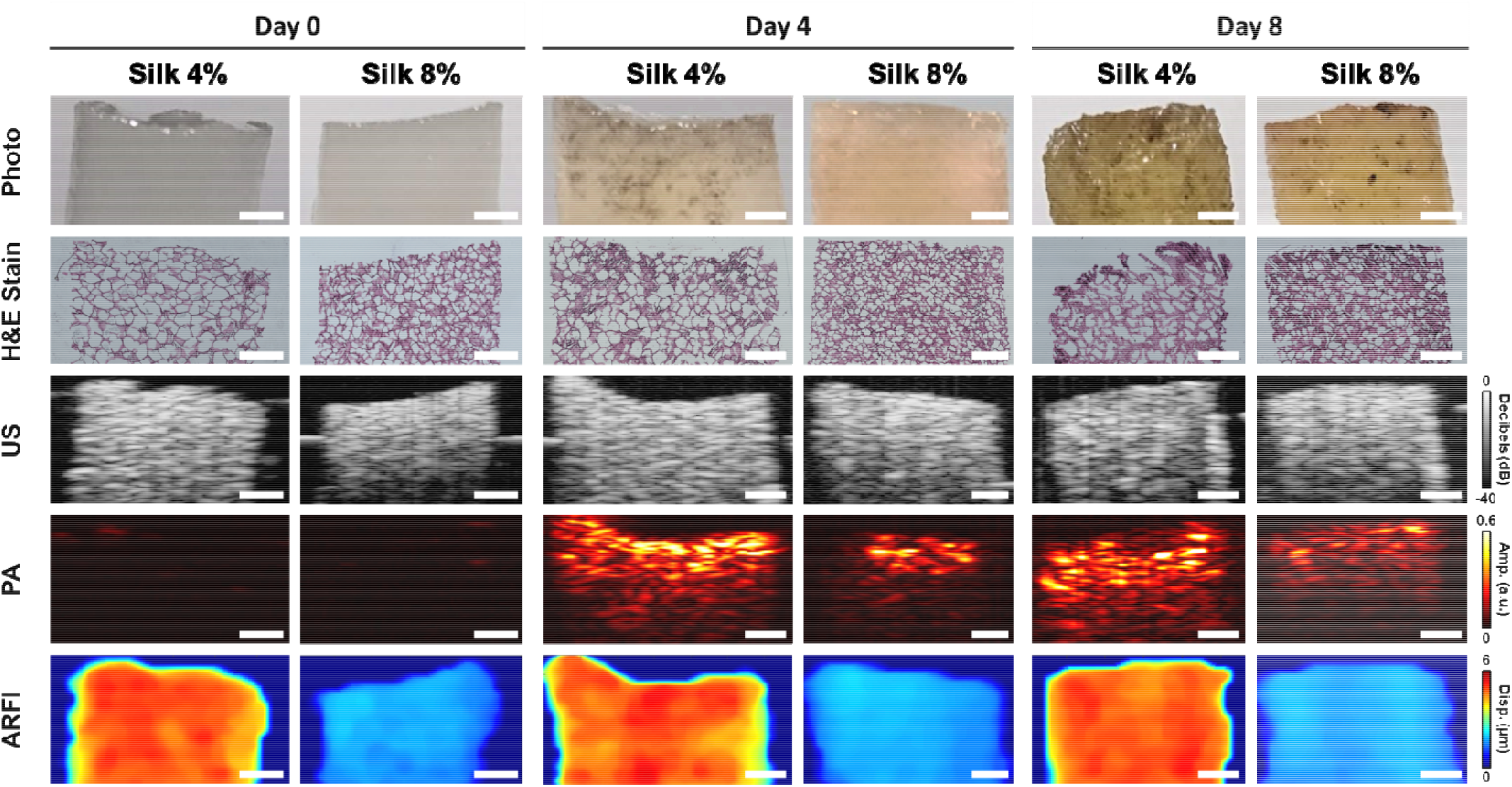
Silk scaffolds seeded with melanoma cells and imaged at different incubation time points. Row 01 displays front facing photos of scaffolds, row 02 displays H&E stained sections of scaffolds, and rows 03 – 05 displays US, PA, and ARFI images, respectively, of scaffolds. All scale bars = 1 mm

When viewing these scaffolds in 3D, 2D observations were maintained as shown in Figure 5 which displays orthogonal slices of day 4 scaffolds. Quantification of these qualitative observations are shown in Figure 6. Figure 6a quantifies the difference in cell growth as a function of silk concentration and incubation duration. Cell growth was defined as the summation of all PA pixel amplitudes above a threshold of 0.03, normalized by the number of US pixels defining the scaffold volume to accommodate for differences in scaffold size. Day 0 pre-seeding expectedly showed little to no PA signal due to the lack of melanoma cells, with any sporadic signal attributed to small air bubbles trapped within the scaffold. After seeding, day 8 expectedly showed statistically more growth than day 4 within both individual silk concentrations. Additionally, it can be observed that between silk concentrations, the 4% silk scaffolds showed more growth than 8% silk for both incubation days as previously mentioned for Figure 4 [28-30].

**Figure 5.**
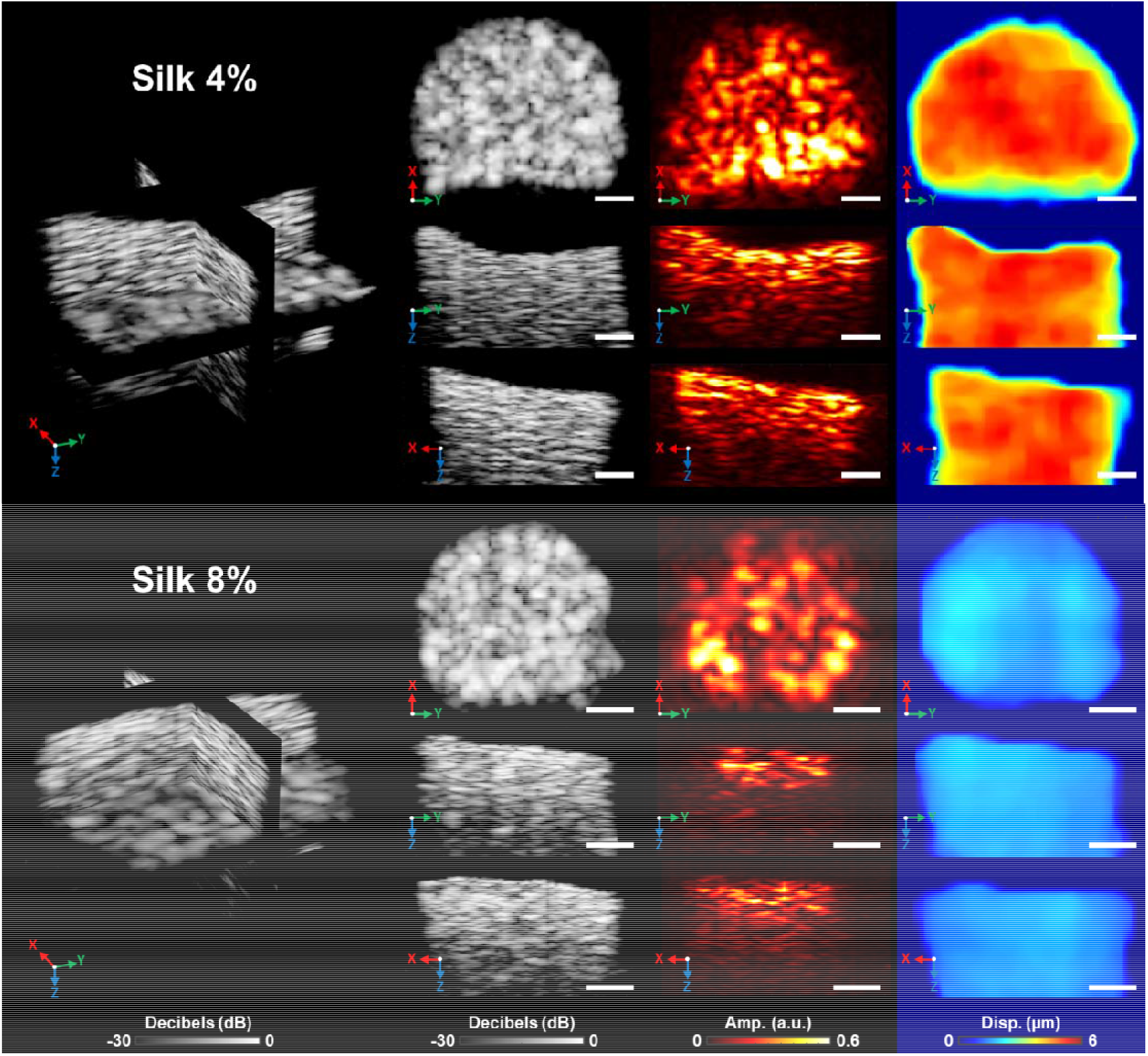
Orthogonal cross sections of 4% and 8% silk on day 04 of incubation to demonstrate 3D visualization of scaffolds. All scale bars = 1 mm

**Figure 6.**
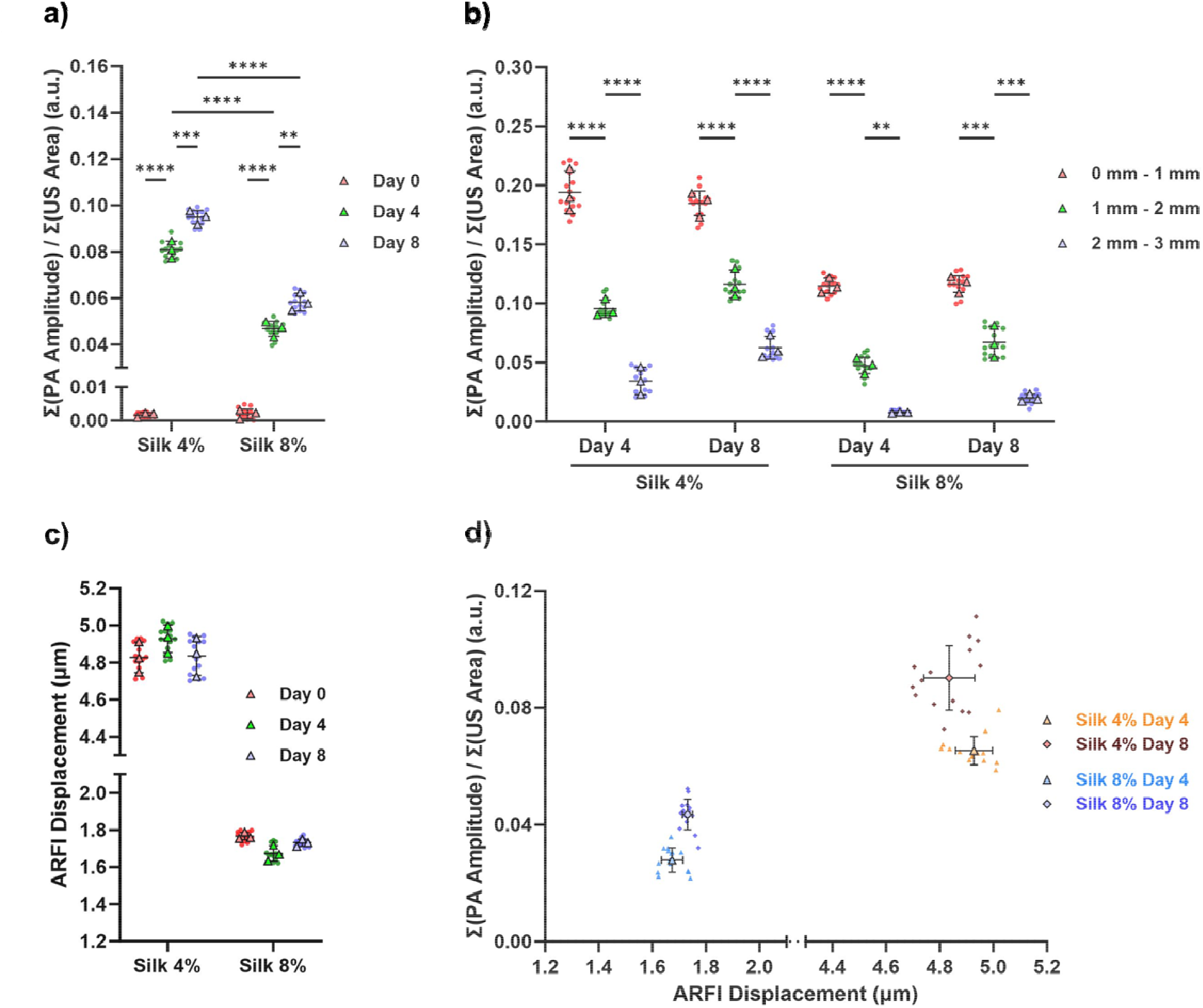
Comparison of seeded scaffolds to determine effects of seeding on scaffold stiffness in addition to scaffold stiffness on proliferation. Outlined data points represent n = 3 scaffolds while non-outlined data points represent individual values averaged for each n. **a)** Average photoacoustic signal across entire scaffold normalized by the corresponding US volume. **b)** Average photoacoustic signal of scaffolds from three different depth ranges normalized by the corresponding US volume. **c)** Average ARFI displacement across incubation days for both silk concentrations. **d)** Comparison between PA amplitude and ARFI displacement. ****p < 0.0001, ***p<0.0005, **p < 0.01. All error bars = s.d.

Following a similar analysis as Figure 6a, Figure 6b quantifies cell proliferation as a function of depth throughout the scaffold. Ultrasound images were used to split the scaffold into three non-overlapping depths at 1 mm thickness each, and within each depth, the PA amplitudes above a threshold of 0.03 were summed and normalized by the number of corresponding US pixels. As expected, the largest amounts of cells were located within the first 1 mm of the scaffold for both days incubation and both silk concentrations as it was the site of seeding. Additionally, at increasing depths, the amount of cells decreased. Figure 6c quantifies the measured ARFI displacement as a function of silk concentration and number of days incubation. For all incubation within individual silk concentrations, displacement values remain relatively consistent all days of ding to the mechanical stability of silk. Note that all statistical differences were confirmed upon performing a two-way ANOVA with a post hoc Tukey multiple comparison test. Figure 6d compares PA amplitude to ARFI displacement to exhibit the relation between cell proliferation and the mechanical stiffness of silk. Both PA amplitude and ARFI displacement were obtained by averaging pixel values within the 1-3 mm depth below the silk surface as Figure 6b confirmed that PA amplitude distinction within the first 1 mm is limited as it was the site of seeding. Thus, differences in cell proliferation and its relation to ARFI displacement will be more apparent below the first 1 mm. From the plot, higher proliferation represented by a higher PA amplitude appears to correlate with softer localized regions of silk represented by higher ARFI displacement. As explained for Figure 4, this may be due to the soft regions characteristically having a lower density of pores resulting in less proliferation resistance.

## 4. Discussion and Conclusions

In this work, ultrasound, photoacoustic, and ARFI were combined into a single microscopy system, referred to as M-PAUR, to provide structural, molecular, and mechanical properties of millimeter-scale tissue engineered constructs. To validate the utility of M-PAUR for tissue engineering, two-part silk scaffolds were constructed and imaged in 3D. Within individual US B-scans, little distinction is observed between the pixel amplitudes located near the top of the two silk concentrations. However, overall structural heterogeneity was observed and easily confirmed visually. Conversely, clear distinctions are observed within PA and ARFI images due to the relatively higher absorption coefficient of graphite compared to silk, and the difference in Young’s modulus of the two silk concentrations. For PA, distinguishing the difference in composition through absorption coefficient demonstrates the microscope’s capability in differentiating the underlying scaffold from other objects. This was further demonstrated through seeding melanoma cells in which PA detected the localized growth of cells. Note that melanoma cells were used due to their high signal contrast from naturally occurring melanin, however, other cell lines may also be detected, provided they display signal contrast either through endogenous chromophores [14], inherently via transfection [31], or through external contrast agents [32, 33]. For ARFI, distinguishing the difference in sample stiffness demonstrates the microscope’s capability in reporting scaffold stiffness and therefore its potential in tracking degradation. As some constructs are designed to be biodegradable and are tuned to specific degradation rates, ARFI provides a method of confirming these characteristics [34]. Additionally, as demonstrated with melanoma seeding, a modest correlation between localized sample stiffness and localized cell growth was observed. Complete correlation was not observed presumably due to variations in the cell seeding process such as location of seeding. However, knowledge of the increased localized stiffness provides the opportunity to homogenize the growth throughout the scaffold by increasing local cell seeding volume.

The current system followed an acoustic-resolution photoacoustic microscopy configuration with an additional transducer for generating an ARFI pulse. Due to the co-linear alignment, this microscope variation can be easily implemented with other photoacoustic microscopes following similar configurations. Furthermore, there is a high probability that this system can be miniaturized and modified to an endoscopic design to track embedded scaffolds [35, 36]. The current system uses a transducer with a central frequency of 20 MHz for imaging and tracking of ARFI displacement. Experimental performance of resolution matched theoretical for photoacoustic and ultrasound, however, ARFI resolution was relatively large which may limit its use for highly inhomogeneous samples. Instead, a higher frequency imaging transducer around 50 MHz in addition to a smaller f/number pushing transducer may be used to lower the ARFI resolution to approximately 200 μm, though the reduction in penetration depth must be considered. Furthermore, the use of 50 MHz will improve US and PA resolution [37]. An additional limitation of our current methodology is the acquisition speed hindered by the low PRF of the OPO laser source. At 10 Hz and an average of 10 shots per A-line for good signal-to-noise (SNR) ratio, large scaffolds may take up to a few hours to fully scan. This may be addressed by introducing an amplifier between the imaging transducer and pulser / receiver to increase the base SNR and allowing for less shots per A-line, or through the application of deep learning to remove noise [38, 39]. A third approach is to use faster sources such as LEDs or laser diodes [40, 41], but they are limited in wavelength bandwidth and still require averaging to obtain good SNR.

Based on the system performance and capability in analyzing the silk scaffolds, M-PAUR and the concept of combining its three synergistic imaging modalities could be applied to a multitude of other applications which would benefit from multi-parametric imaging. Our future work includes investigating the capability of imaging multiple cell lines by leveraging various photoacoustic genetic probes (spectroscopic imaging), and imaging vascular profiles within the 3D scaffolds. As the field of tissue engineering continues to advance to fabricate larger tissues and organs, the importance of non-ionizing and non-invasive imaging systems, particularly at meso-scale, becomes increasingly apparent. By facilitating a deeper understanding and better characterization of scaffold architectures and cellular proliferation, such imaging systems will be pivotal in driving the success of future tissue-engineering solutions.

## Acknowledgements

The authors would like to acknowledge support from Tufts School of Engineering, Tufts Clinical and Translational Science Institute Pilot award (Mallidi) and P41EB027062 (Kaplan). The authors would like to thank Charles Swan and Annika Schaad for help in constructing the custom fiber bundle and Aayush Arora for help in cell culturing.

## Conflicts of Interest

The authors declare that they have no known competing financial interests or personal relationships that could have appeared to influence the work reported in this paper.

## Notes

### Competing Interest Statement

The authors have declared no competing interest.

