## Supplemental Data for "Multi-spectral photoacoustic imaging combined with acoustic radiation force impulse imaging for applications in tissue engineering"

**Supplementary Figure S1**

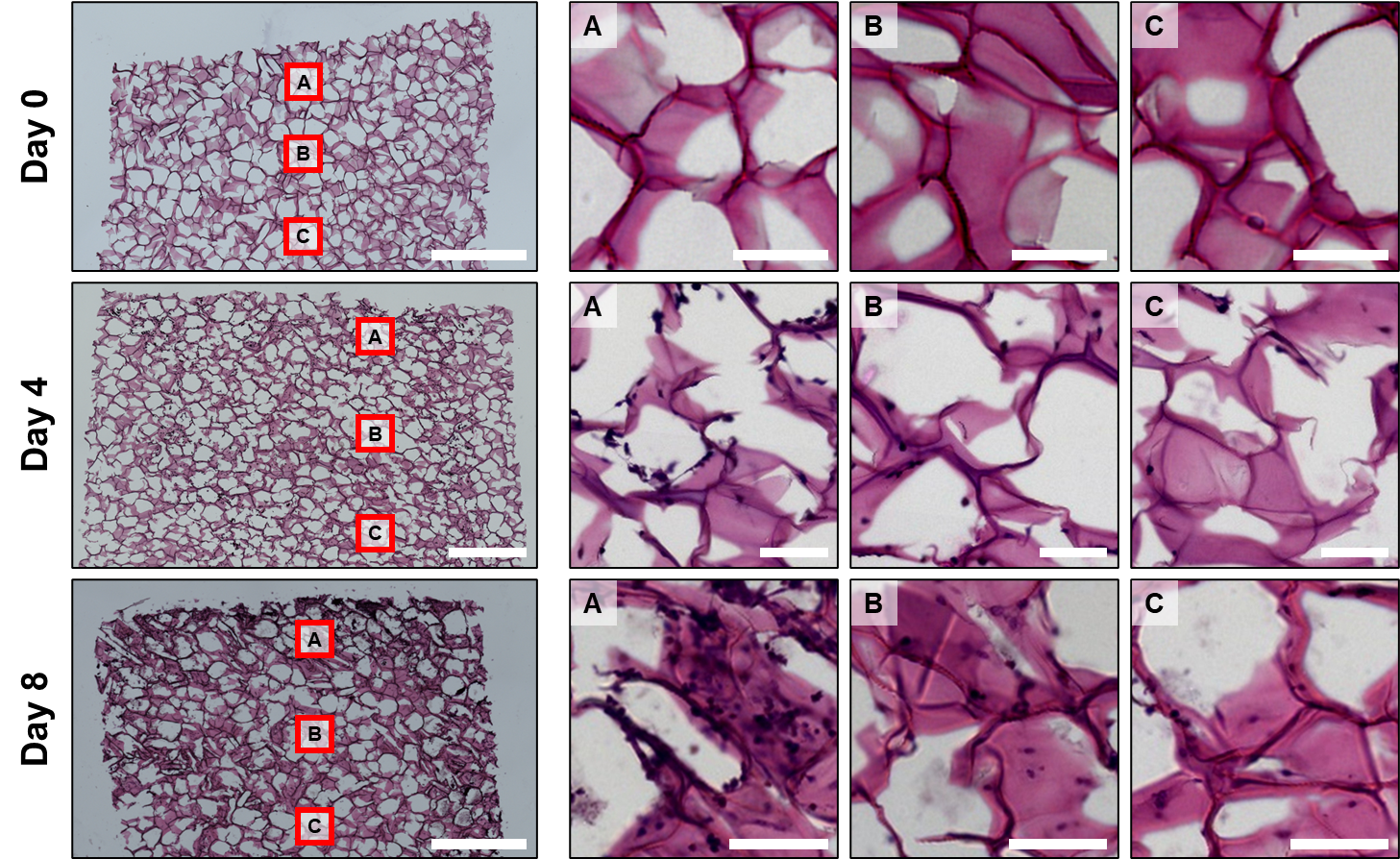

Figure S1. Histology stains of 8% silk for progressive days of incubation. Left column shows full slide images. Scale bar = 1 mm. Subsequent columns correspond to image insets starting from top (A) to bottom (C). Scale bar = 100 µm.
